# Predicting Molecular Docking Affinity of Per- and Polyfluoroalkyl Substances (PFAs) Towards Human Blood Proteins Using Generative AI Algorithm DiffDock

**DOI:** 10.1101/2023.08.03.551898

**Authors:** Dhan Lord B. Fortela, Ashley P. Mikolajczyk, Miranda R. Carnes, Wayne Sharp, Emmanuel Revellame, Rafael Hernandez, William Holmes, Mark Zappi

## Abstract

This study computationally evaluates the molecular docking affinity of various perfluoroalkyl and polyfluoroalkyl substances (PFAs) using a generative machine learning algorithm, DiffDock, specialized in protein-ligand blind-docking learning and prediction. Concerns about the chemical pathways and accumulation of PFAs in the environment and eventually in human body has been rising due to empirical findings that levels of PFAs in human blood has been rising. Though there is currently a heightened need to understand the pathways of PFAs, empirical studies on PFAs have been relatively slow due to the time-scale and cost of standard chemical analysis such as those in blood samples. The current study demonstrates the implementation of DiffDock and assesses the prediction results in relation to empirical findings. The capability of an advanced generative artificial intelligence (AI) algorithm designed for protein-ligand docking such as DiffDock offers a fast approach in determining the potential molecular pathways of PFAs in human body.

## 1. Introduction

The risks to human health and environment of per- and polyfluoroalkyl substances (PFAs) have been a major concern in the current decade due to their prevalence in water, soil, air, and food [1, 2]. PFAs have been used in industrial manufacturing and in consumer products due to their useful properties [3]. Current scientific research suggests that exposure to certain PFAs may be harmful to human health [4, 5]. This growing concern about the environmental and anthropologic pathways of PFAs has prompted health and environmental regulatory agencies such as the US National Institutes of Health (NIH) [6] and US Environmental Protection Agency (EPA) to implement strategic plans to address PFAs contamination and accumulation [1]. The US Centers for Disease Control and Prevention (CDC) has recently established blood testing procedures for PFAs as part of the efforts to understand PFAs health effects [7]. A chemical analysis of human blood plasma indicates that albumin is the major carrier protein for PFAs [8]. Though such blood chemistry analyses are currently being developed and tested via case studies [9], human blood is a complex matrix of biomolecules [10] in which established chemical analysis techniques may not be able to easily detect PFAs complexed with blood components. With human blood serving many functions including the critical task of distributing nutrients to various parts of the body [11], blood-based analyses provides critical insights about human health [12] such as risks with PFAs.

A potential powerful tool that can be used to determine the molecular docking affinity of PFAs with human blood proteins is the generative artificial intelligence (AI) algorithm called DiffDock [13], which is the latest significantly improved generative-learning algorithm for molecular docking of a ligand (or a small molecule) to protein receptor trained on large dataset of protein-ligand complexes [14]. Though originally intended for drug-discovery applications, DiffDock can be applied to other similar protein-ligand docking problems because DiffDock was trained by Corso, Stärk [13] on 17,000 protein-ligand complexes from the Protein Data Bank (PDB) [15]. DiffDock learns over the manifold of ligand poses consists of translational, rotational, and torsional dimensions by implementing a diffusion generative model [13, 16].

This work implements the DiffDock algorithm in the prediction of molecular docking of PFAs with blood proteins to illustrate potential uses outside of its original intended application in drug-discovery. This work is a first demonstration on the use of generative AI in estimating the affinity of PFAs to bind onto human blood proteins. Even though the work is limited to select human blood proteins, the findings of the work may usher further implementations of a generative AI algorithm such as DiffDock in elucidating molecular binding of PFAs onto other proteins in the human body. In a more general view, the use of generative AI in molecular docking prediction may aide in fast and comprehensive studies of the pathways of PFAs into human bodies and other living organisms, and in the improved design of materials, processes, and technologies to minimize or eliminate bioaccumulation of PFAs and other contaminants.

## 2. Methodology

A schematic of the data analytics workflow implemented is shown in Figure 1.

**Figure 1.**
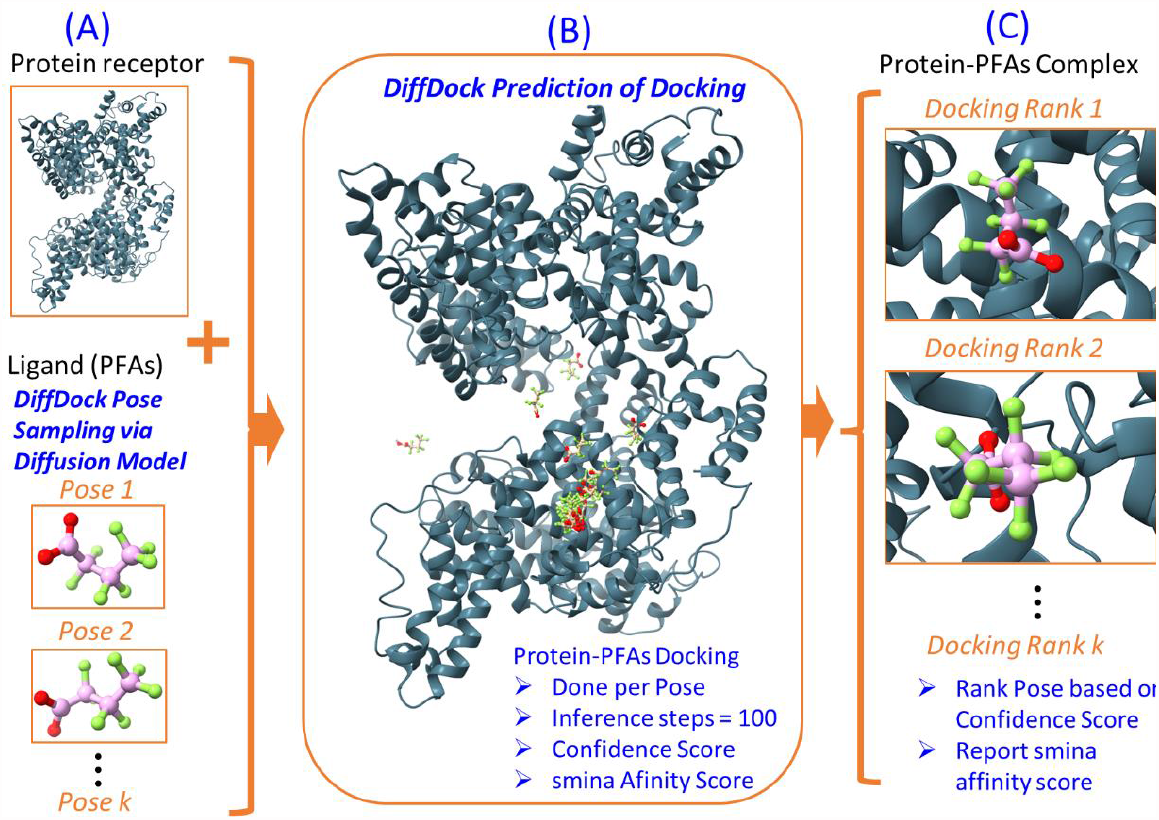
Schematic overview of the data analytics workflow implemented in this study. The example docking shown is that of protein receptor albumin (1AO6) and ligand perfluorobutanoic acid (PFBA). (A) The conformation of the ligand molecule is sampled via diffusion generative model in DiffDock resulting to a ligand pose (with all information about atomic positions in the molecule) to be docked to the protein receptor. (B) The DiffDock algorithm takes the ligand pose and docks the ligand to the protein by searching for the docking position of higher likelihood with 100 inference steps using the probability distribution developed in the trained DiffDock model. The convergence of PFBA to a certain location in the albumin protein is apparent in the Figure 1B. The associated confidence score and smina affinity score of the docking is also computed. Then another ligand pose is generated, and the docking computations are repeated until all target ligand poses are docked. (C) The confidence score of docking by each ligand pose is then used to rank the ligand poses, and the smina affinity scores are reported. A video clip of the docking inference steps of PFBA-albumin to convergence of Rank 1 pose is provided as supplementary material and in the GitHub repository of the paper.

The DiffDock algorithm computations were implemented using Python coding language [17, 18] run in GoogleColab using NVIDIA Tesla T4 GPU runtime [19]. Each run of protein-ligand docking computations took 30 minutes on the average to complete, hence, all runs of 42 protein-ligand pairs resulted to a total of 1260 minutes (21 hours) of DiffDock runs for the whole dataset used in this paper. A copy of the Python codes used are provided as Jupyter Notebook [20] file in the GitHub repository for the paper: https://github.com/dhanfort/Blood_Protein_PFAs_Docking.git [21]. The 3D renderings of protein and molecular structures were done using the ChimeraX software [22, 23]. The structural formula of PFAs were rendered using the OpenBabel software [24].

### 2.1 Study Dataset: Blood Proteins and PFAs Molecular Structures

The human blood proteins used in this analysis were selected to represent key functions in blood serum: (1) albumin, which is 55% of the blood plasma proteins [25], (2) hemoglobin, which is the transport protein in red blood cells [26], (3) alpha-1-antitrypsin [27], which is a serpin globulin that is typically concentrated from donated bloods and used for therapy of certain disorders, and (4) corticosteroid-binding globulin (CBG), which is a glycoprotein that binds to cortisol and other glucocorticoids in the blood and helps with anti-inflammatory actions [28]. The blood proteins structures format used was the “.pdb” and the protein molecular data were downloaded from the Protein Data Bank (PDB) online repository [15] via PDB identification label of each protein: 1AO6 for albumin [29, 30], 1A3N for hemoglobin [31, 32], 1QLP for alpha-1-antitrypsin [33, 34], and 2VDY for corticosteroid-binding globulin (CBG) [35, 36]. A 3D rendering of the protein structures are shown in Figure 2.

**Figure 2.**
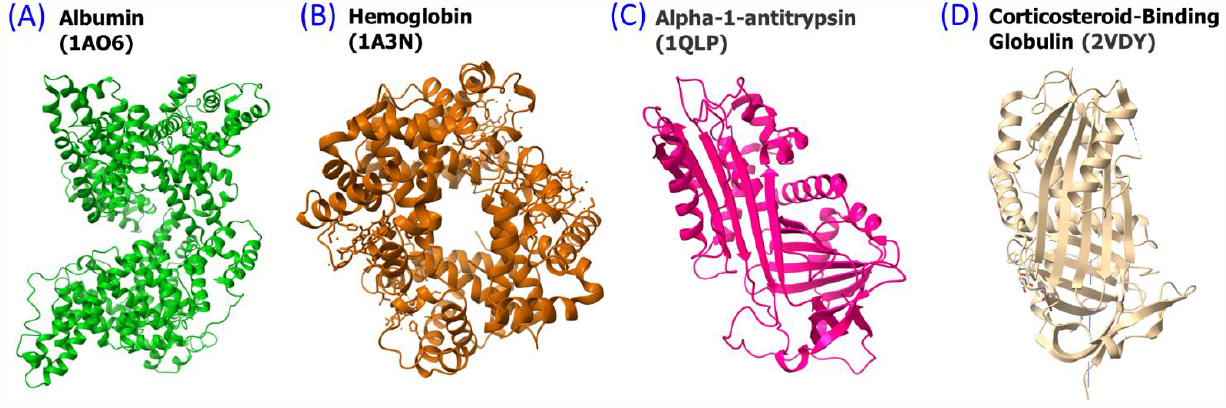
A 3D structure rendering of the human blood proteins used in this work: (A) Albumin with PDB identification number 1AO6, (B) Hemoglobin with PDB identification number 1A3N, (C) Alpha-1-antitrypsin with PDB identification number 1QLP, and (D) Corticosteroid-binding globulin (CBG) with PDB identification number 2VDY.

The PFAs molecular structures format used was the simplified molecular-input line-entry system (SMILES) [37] downloaded from PubChem [38] as shown in Table 1. The selection of the PFAs was based on the result of review of current literature on PFAs with focus on the reports of US EPA [1], NIH [6], and published empirical analysis of PFAs in human blood [8, 39, 40]. Hence, there are 12 PFAs used in the current work ranging from short-chain to long-chain structures (Table 1). For a comprehensive analysis, the structural formula and the 3D ball-and-stick rendering of the PFAs are also presented in Figure 3. The PFAs are as follows: perfluorobutanoate (PFB), perfluorobutanoic acid (PFBA), hexafluoropropylene oxide dimer acid (HFPO-DA), perfluorooctanesulfonic acid (PFOS), perfluorooctanoic acid (PFOA), perfluorononanoic acid (PFNA), perfluorohexanesulfonic acid (PFHxS), perfluorobutanesulfonic acid (PFBS), perfluoropentanoic acid (PFPeA), perfluorohexanoic acid (PFHxA), perfluoroheptanoic acid (PFHpA), and perfluorodecanoic acid (PFDA).

**Table 1.**
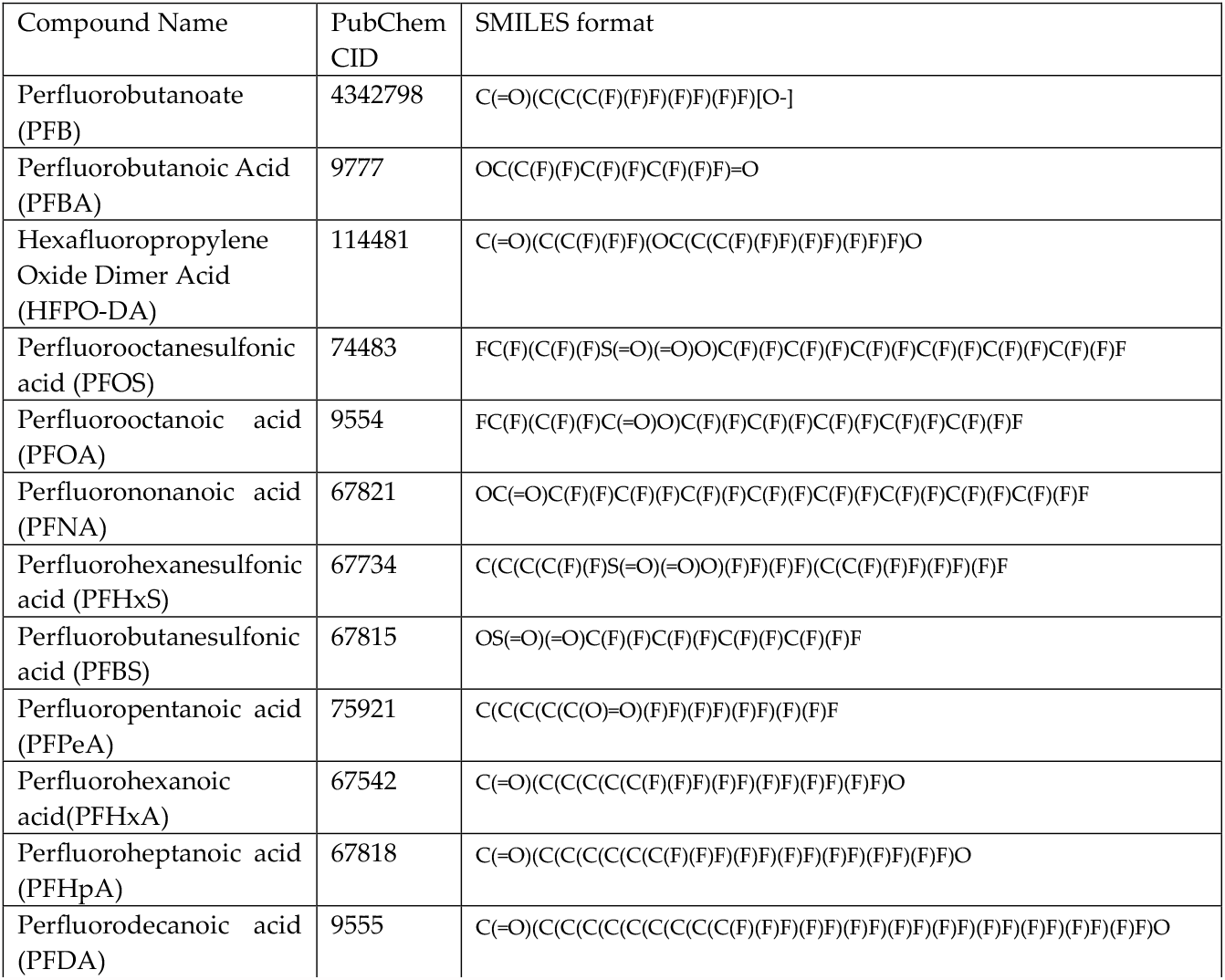
PFAs used in this study with their respective PubChem CID identification number and SMILES format.

**Figure 3.**
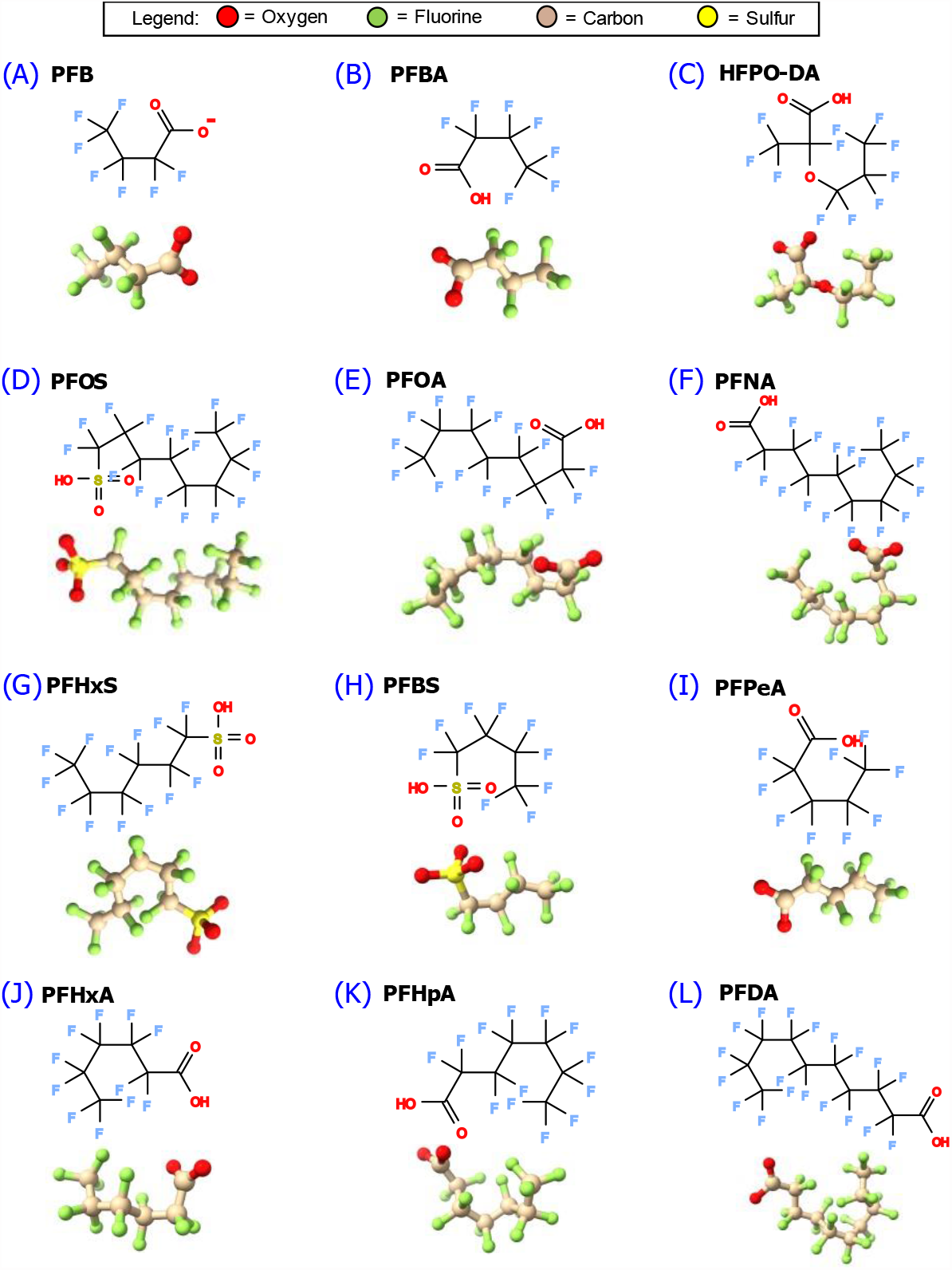
Structural formula and ball-and-stick rendering of the PFAs molecules used in the molecular docking computations with blood proteins: (A) PFB, (B) PFBA, (C) HFPO-DA, (D) PFOS, (E) PFOA, (F) PFNA, (G) PFHxS, (H) PFBS, (I) PFPeA, (J) PFHxA, (K) PFHpA, (L) PFDA.

### 2.2 DiffDock: Docking Inference, Docking Rank, and Binding Affinity Score

The DiffDock algorithm implements a generative diffusion model (a.k.a., Brownian motion) during training and inference steps [13]. This generative task allows the efficient sampling of translational, rotational, and torsional parameters of a ligand as it docks with the protein receptor [13]. Since DiffDock has been trained on a large dataset of protein-ligand complexes (17,000 protein-ligand complexes from PDB; see Corso, Stärk [13] for the details), the pre-trained model can be used for inference of protein-ligand complex systems. The goodness-of-fit of protein-ligand docking can be measured using the proposed metrics by the originators of the algorithm [13]: (1) Rank of docking based on Confidence Score, and (2) smina Affinity Score. These metrics used in this study are discussed as follows.

#### 2.2.1 Molecular Docking Rank

The molecular docking pose rank (Equation 1) was based on the Confidence Score (Equation 2) as proposed by Corso, Stärk [13], which was based on the concept by Song, Sohl-Dickstein [16]. The pertinent equations implemented in the algorithm for this docking rank are as follows. The rank of protein-ligand docking pose is evaluated in DiffDock by sorting the Confidence Score (Equation 2), where Rank 1 is the docking with the highest Confidence Score.

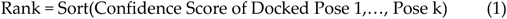

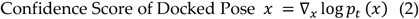

Equation 2 describes the score of each marginal distribution of ligand pose *x*, which is the assignment of atomic positions in ℝ^3^ (3-dimensions) [13], at each time step *t* of the generative diffusion process [16]. Hence, a ligand with *n* atoms will have ℝ^3*n*^ dimensions in every pose *x*. The gradient ∇_*x*_ is the result of framing the diffusion process as stochastic differential equations (SDEs) according to Song, Sohl-Dickstein [16], and *p* denotes the ligand pose probabilistic distribution. Rigorous derivations of Equation 1 and 2 are presented by Corso, Stärk [13] and Song, Sohl-Dickstein [16].

#### 2.2.2 smina Affinity Score

The scoring and minimization with AutoDock Vina (smina) affinity score by Koes, Baumgartner [41] was also integrated for a comprehensive evaluation of the correct docking poses. The general form of the equation implemented in the algorithm for the smina affinity score computations is shown in Equation 3, where *c* is the smina affinity score (in kcal/mol). The smina score can be positive or negative depending on the strength of the interaction between the protein and the ligand. A negative smina score indicates a stronger binding affinity while a positive smina score indicates weaker binding affinity [41]. The smina affinity score is a measure of the standard chemical potential of the protein-ligand system [42]. Hence, the smina affinity score is an approximation of the Gibbs free energy of binding between the protein receptor and a ligand [43]. The smina affinity score takes the unit of energy per mole, specifically kcal/mol in the DiffDock algorithm [13], which is consistent with AutoDock Vina [42]. The chemical potential of the protein-ligand complex is minimized when the smina affinity score is minimized [42]. Equation 3 is essentially the sum of intermolecular forces *c*_*inter*_ and intramolecular forces *c*_*intra*_ [42].

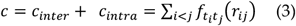

The summation in Equation 3 is over all pairs of atoms that can move relative to each other. The type of atom *i* is designated by a type *t*_*i*_ and a set of interactions functions 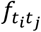 with the interatomic distance *r*_*ij*_ [42]. The details of the various components of Equation 3 have been published elsewhere [41-43]. The computation of smina affinity score have been included in the Python codes used to run the DiffDock algorithm and are part of the Jupyter Notebook file in the paper online repository [21].

## 3. Results and Discussion

The results of DiffDock inferences on the molecular docking of PFAs with blood proteins are presented in Figures 4, 5, 6 and 7. In these, the pose docking ranks are plotted against the smina affinity scores. The Top-1 docking smina affinity scores are presented in Table 2, and the average of the Top-5 docking smina affinity scores are presented in Table 3.

**Table 2.** Summary of smina affinity scores (kcal/mol) of the Top-1 docking pose of PFAs towards the blood proteins.

| PFA Compound | Albumin | Hemoglobin | Alpha-1-antitrypsin | Corticosteroid-binding globulin (CBG) |
| --- | --- | --- | --- | --- |
| PFB | -2.72 | 0.58 | 13.06 | -2.54 |
| PFBA | -2.34 | -3.59 | 7.49 | -2.63 |
| HFPO-DA | -1.90 | 0.26 | 29.69 | -5.13 |
| PFOS | 2.83 | 3.41 | 58.60 | -6.33 |
| PFOA | -3.90 | 6.36 | 39.43 | -4.33 |
| PFNA | -1.48 | 5.82 | 31.85 | -5.13 |
| PFHxS | -4.88 | 7.41 | 27.76 | -4.39 |
| PFBS | -3.54 | 3.74 | 19.99 | -5.35 |
| PFPeA | -3.78 | -1.80 | 9.96 | -5.05 |
| PFHxA | -2.50 | 3.42 | 17.47 | -3.88 |
| PFHpA | -4.07 | 2.41 | 26.34 | -5.92 |
| PFDA | 1.97 | 6.64 | 64.53 | -2.10 |

**Table 3.** Summary of average smina affinity scores (kcal/mol) of the Top-5 docking poses of PFAs towards the blood proteins. Numbers inside parentheses are sample standard deviations.

| PFA Compound | Albumin | Hemoglobin | Alpha-1-antitrypsin | Corticosteroid-binding globulin (CBG) |
| --- | --- | --- | --- | --- |
| PFB | -2.33 (1.45) | 1.24 (1.53) | 10.89 (2.58) | -2.86 (0.36) |
| PFBA | -2.46 (1.03) | -1.44 (2.23) | 12.11 (2.91) | -2.23 (0.60) |
| HFPO-DA | -2.74 (1.08) | 5.98 (5.64) | 29.80 (5.70) | -4.31 (0.84) |
| PFOS | -0.38 (2.37) | 5.39 (2.27) | 65.17 (15.71) | -4.13 (1.71) |
| PFOA | -2.71 (0.88) | 5.77 (0.76) | 47.43 (6.95) | -4.87 (0.48) |
| PFNA | -1.76 (1.43) | 5.26 (0.94) | 53.40 (19.53) | -4.57 (0.46) |
| PFHxS | -2.28 (1.65) | 7.54 (1.52) | 42.20 (10.11) | -3.71 (1.27) |
| PFBS | -2.37 (0.83) | 5.36 (1.68) | 22.66 (3.06) | -4.25 (0.77) |
| PFPeA | -2.97 (0.52) | -0.52 (1.55) | 16.21 (5.10) | -4.40 (0.83) |
| PFHxA | -2.51 (0.36) | 3.38 (1.78) | 19.77 (4.25) | -3.88 (0.80) |
| PFHpA | -2.46 (1.11) | 2.42 (1.24) | 32.36 (11.02) | -5.47 (0.58) |
| PFDA | 3.05 (3.55) | 7.99 (2.16) | 67.16 (7.60) | -3.88 (1.23) |

**Figure 4.**
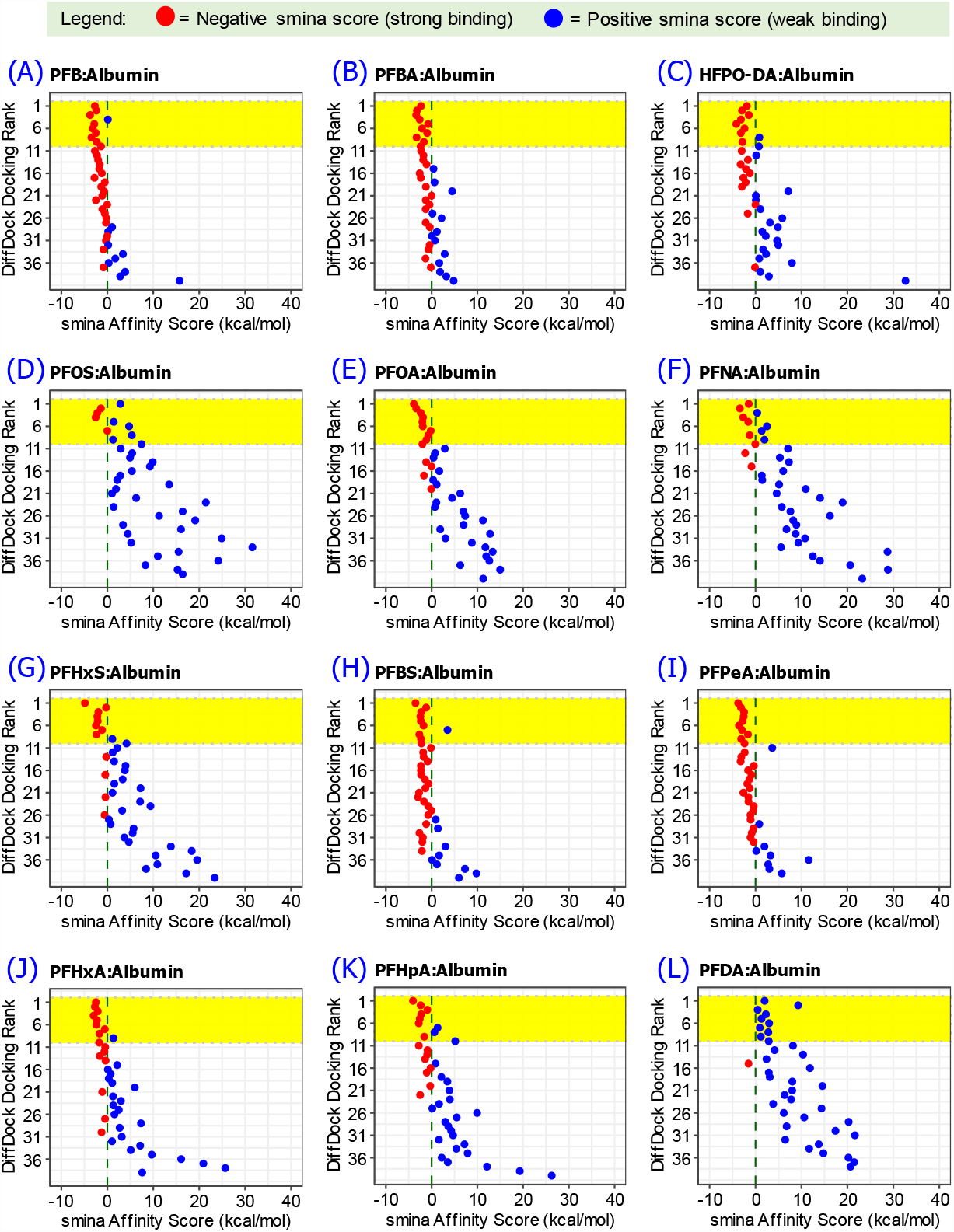
Molecular docking pose rank computed using DiffDock and the associated smina affinity score between albumin protein molecule and PFAs. Top-10 docking ranks are highlighted in yellow. (A) PFB, (B) PFBA, (C) HFPO-DA, (D) PFOS, (E) PFOA, (F) PFNA, (G) PFHxS, (H) PFBS, (I) PFPeA, (J) PFHxA, (K) PFHpA, (L) PFDA.

**Figure 5.**
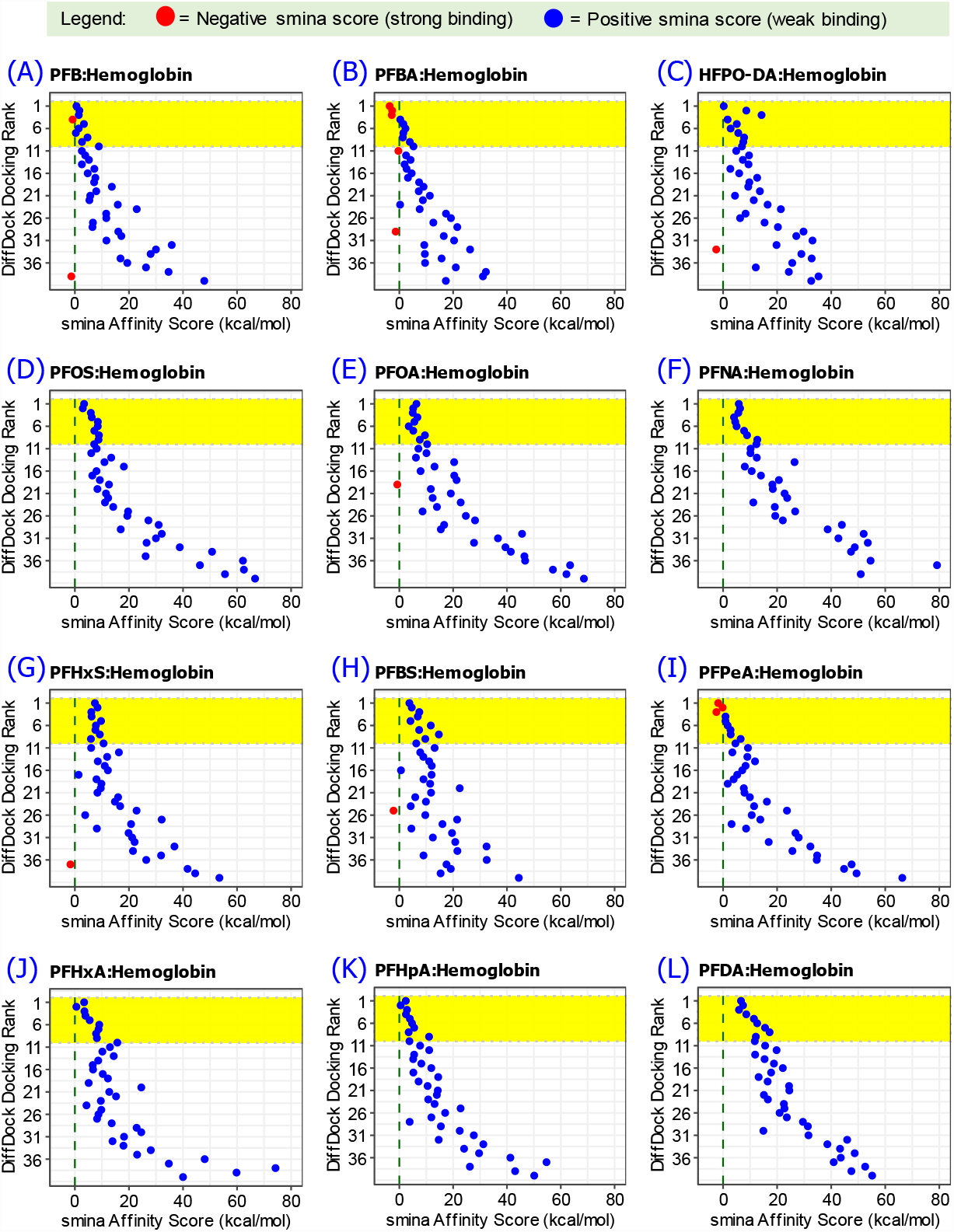
Molecular docking pose rank computed using DiffDock and the associated smina affinity score between hemoglobin protein molecule and PFAs. Top-10 docking ranks are highlighted in yellow. (A) PFB, (B) PFBA, (C) HFPO-DA, (D) PFOS, (E) PFOA, (F) PFNA, (G) PFHxS, (H) PFBS, (I) PFPeA, (J) PFHxA, (K) PFHpA, (L) PFDA.

**Figure 6.**
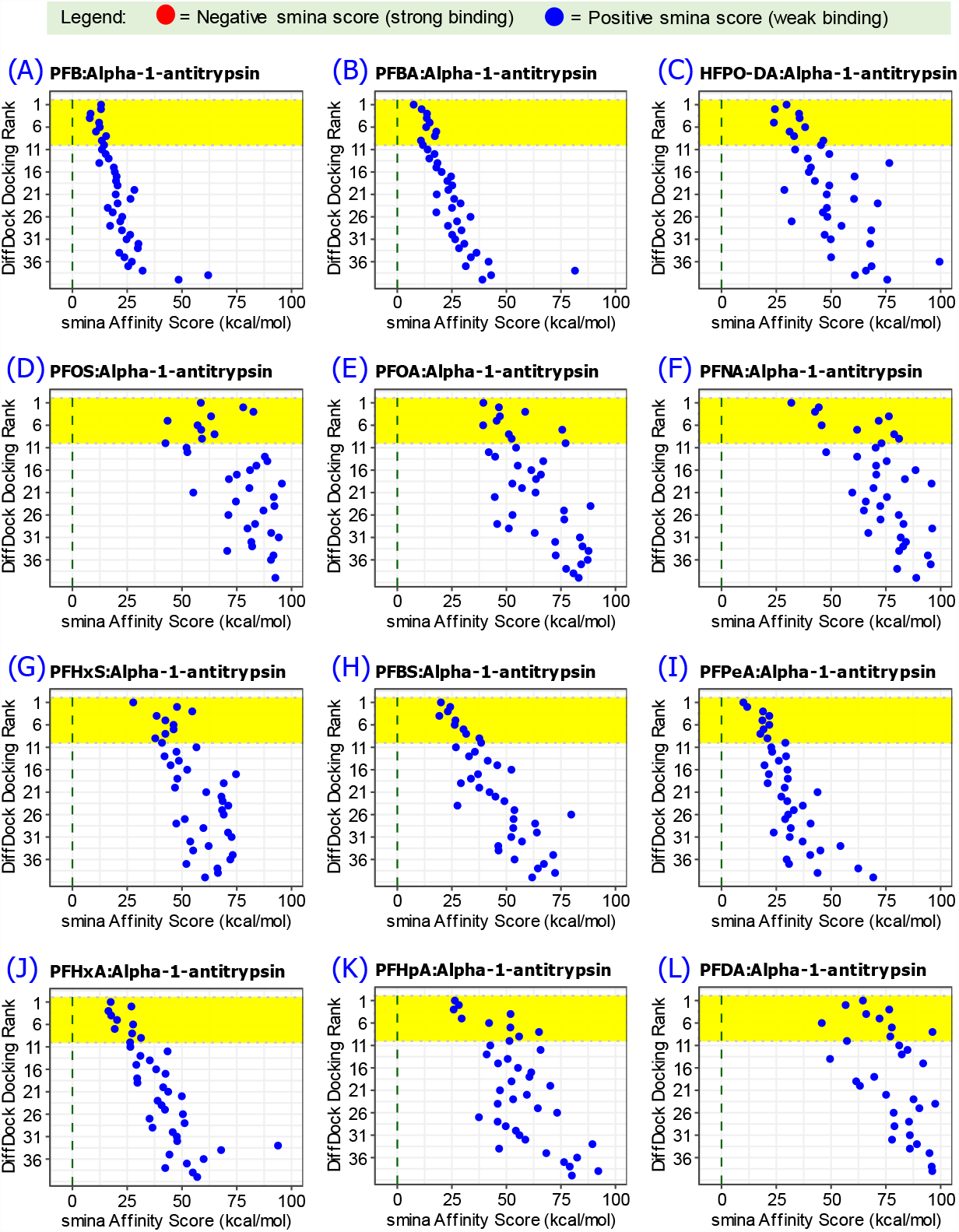
Molecular docking pose rank computed using DiffDock and the associated smina affinity score between alpha-1-antitrypsin protein molecule and PFAs. Top-10 docking ranks are highlighted in yellow. (A) PFB, (B) PFBA, (C) HFPO-DA, (D) PFOS, (E) PFOA, (F) PFNA, (G) PFHxS, (H) PFBS, (I) PFPeA, (J) PFHxA, (K) PFHpA, (L) PFDA.

**Figure 7.**
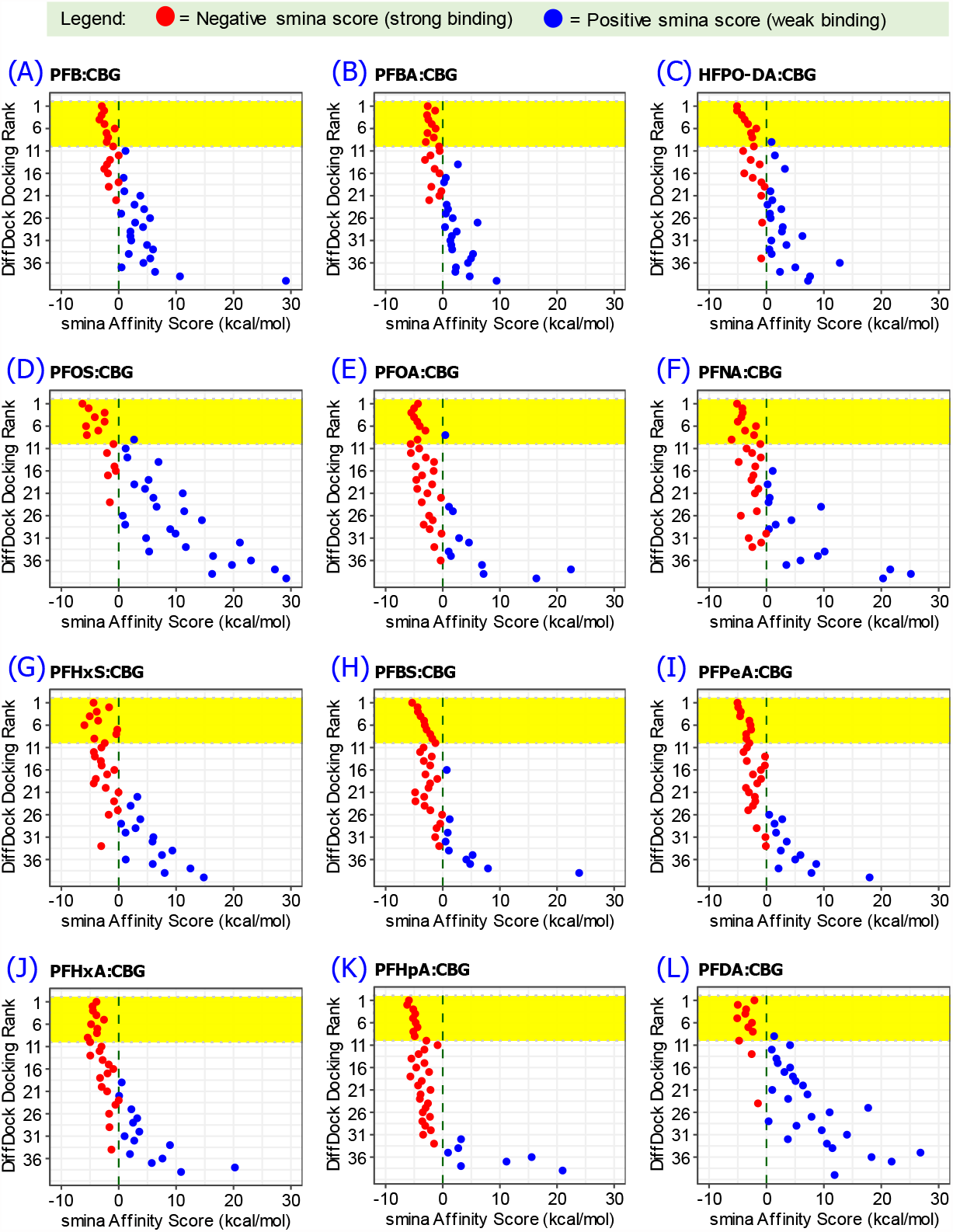
Molecular docking pose rank computed using DiffDock and the associated smina affinity score between corticosteroid-binding globulin (CBG) protein molecule and PFAs. Top-10 docking ranks are highlighted in yellow. (A) PFB, (B) PFBA, (C) HFPO-DA, (D) PFOS, (E) PFOA, (F) PFNA, (G) PFHxS, (H) PFBS, (I) PFPeA, (J) PFHxA, (K) PFHpA, (L) PFDA.

### 3.1. Trends of Docking Rank and Binding Affinity

The plots of DiffDock protein-PFAs docking ranks versus smina affinity scores (Figure 4, 5, 6 and 7) show a general trend of the smina affinity scores becoming more negative as the docking improves with the top-rank docks having the top confidence scores. Because the smina affinity score is an approximation of the Gibbs free energy of the system [43], the more negative the smina affinity score becomes, the more stable the protein-PFAs complex being formed. Note that the smina affinity score was calculated after the docking inference steps, which implies that the trends exhibited in the rank-versus-affinity score plots (Figure 4, 5, 6 and 7) confirm how the diffusion generative model of DiffDock guides the inference of ligand pose and docking to a state consistent with the principles of molecular thermodynamics where a chemical system attains stability by seeking very negative Gibbs free energy [44].

The smina affinity score (kcal/mol) with the most negative values are close to the range of published experimental and simulation-based binding free energy of stable protein-ligand complex at around -8 kcal/mol [45]. The apparent randomness in the smina affinity scores in the sequence of the docking ranks is due to the probabilistic approach of the diffusion generative model in DiffDock sampling the ligand pose [13]. The convergence to optimal docking position results in the general trend of favoring more negative smina score as docking rank improves.

### 3.2. Comparison with Related PFAs Empirical Results

It can be observed that many of the PFAs have very negative smina scores towards albumin and CBG, which indicates strong binding affinity (Table 2 and 3). On the other hand, almost all of the PFAs have weak binding affinity towards hemoglobin and alpha-1-antitrypsin (Table 2 and 3). Based on top-1 docking, PFHxS among all the PFAs has the strongest binding towards albumin with smina score of -4.88 kcal/mol, and PFOS among all the PFAs has the strongest binding towards CBG with smina score of -6.88 kcal/mol.

Though studies have shown that glycated hemoglobin (HbA1c) has some positive correlation with PFOA concentration in blood [46, 47], there has not been definitive reports of complex formation between hemoglobin and PFAs. The trends of PFAs-hemoglobin complex docking shown in Figure 5 implies very weak binding affinity between these protein-ligand systems. This suggests that the determined correlation between PFAs and HbA1c may be the result of indirect mechanistic association.

The selection of alpha-1-antitrypsin for this study is because of the lack of studies on its chemistry with PFAs amid the fact that it has been recognized as a glycoprotein that may have multifunctional roles with therapeutic effects in many inflammatory and autoimmune diseases [48]. With this value, the potential of alpha-1-antitrypsin as carrier of PFAs to other individuals via blood transfusion must be evaluated. The results in Figure 6 show that all PFAs tested have very weak binding affinity towards alpha-1-antitrypsin protein as indicated by the positive smina scores. This may mean that the risk of carrying PFAs from one person to another with alpha-1-antitrypsin from blood is very low.

### 3.3. Significance of the Current Work

Reviews on the literature of PFAs have found that more than 4700 of these substances have been introduced into many of the products and ecosystems that are used in everyday life [49-51]. This large pool of PFAs poses an inherent challenge in conducting studies on their pathways in the environment and in human body [49]. However, the closely related field of drug-discovery has been recently making significant progress in solving the same problem of studying numerous possible compounds that may affect the human body [13]. One of the common steps in both study areas is the analysis of molecular docking of protein-ligand complexes. Hence, this work demonstrates the potential to accelerate analysis of PFAs affinity to human body using the generative AI algorithm DiffDock, which originated as a computation tool in drug-discovery [13].

On top of screening the strong and week binding of protein-PFAs complexes, the use of a computational algorithm like DiffDock may also help elucidate chemical pathways. An example case is that of PFOS, which has been reported by empirical measurements that it strongly complexes with albumin [8]. The DiffDock computation results do not suggest this finding (Figure 4D), and show that other proteins may be responsible for the strong binding of PFOS to blood component such as CBG as shown in Figure 7D and Table 2. With this intriguing result, the relevant literature of PFOS measurements in human blood was then reviewed and it was found that these measurements involve the physical separation of blood components before PFOS and other PFAs are measured [8]. The fractionation of human blood via ultracentrifuge used by Forsthuber, Kaiser [8] can reach only 85-90% purity for albumin fraction [52]. This leads to the possibility that other proteins such as CBG are not completely separated from albumin and are the proteins carrying PFOS via complex formation (Figure 7D) and not albumin (Figure 4D). Another method used in blood fractionation is electrophoresis, but this method is not selective towards albumin, which migrates toward the anode but at slower rate compared to the prealbumin proteins transthyretin (sometimes called thyroxine-binding prealbumin) and the retinol-binding protein [53]. Hence, the current methods of separating albumin from the rest of the blood components may affect the measurements on whether PFAs are complexed with albumin. This analysis of PFOS trends illustrates the kind of investigation that follows after molecular docking analysis is performed and can lead to more precise measurements of PFAs complexes and their pathways in the human body.

## 4. Conclusions

Generative AI algorithm DiffDock accelerates protein-PFAs molecular docking computations. The various PFAs exhibit varied levels of binding affinity towards human blood proteins. Of the four human blood proteins, albumin and corticosteroid-binding globulin (CBG) results to the most negative smina affinity scores indicating strong protein-PFAs binding while PFAS complex with hemoglobin and alpha-1-antitrypsin are weak. PFOS exhibits weak binding with albumin, and strong binding affinity towards CBG.

## Supplementary Materials

The following supporting information are available in the GitHub repository of the paper: Video S1: DiffDock inference on PFBA with albumin, and Video S1: DiffDock inference on PFOS with alpha-1-antitrypsin.

## Author Contributions

Conceptualization, D.L.B.F., A.P.M. and W.S.; methodology, D.L.B.F., A.P.M. and M.R.C.; software, D.L.B.F. and M.R.C.; formal analysis, D.L.B.F., A.P.M., W.S., E.R., R.H., W.H. and M.Z.; data curation, D.L.B.F.; project administration, D.L.B.F.; funding acquisition, D.L.B.F., M.R.C. and M.Z. All authors have read and agreed to the published version of the manuscript.

## Data Availability Statement

The Python codes used in implementing the DiffDock algorithm were based on the codes originally developed by Corso, Stärk [13]. The codes are made available as Jupyter Notebook files in the online repository of the paper via GitHub: https://github.com/dhanfort/Blood_Protein_PFAs_Docking.git [21].

## Acknowledgments

We thank the supportive staff and students of the Department of Chemical Engineering and the Energy Institute of Louisiana at the University of Louisiana at Lafayette. We are also grateful to LaSPACE for supporting students who pursue research in STEM projects.

